# Nociceptor subtypes are born continuously over DRG development peaking at E10.5—E11.5

**DOI:** 10.1101/2021.07.26.453909

**Authors:** Mark A. Landy, Megan Goyal, Helen C. Lai

## Abstract

Sensory neurogenesis in the dorsal root ganglion (DRG) occurs in two waves of differentiation with larger, myelinated proprioceptive and low-threshold mechanoreceptor (LTMR) neurons differentiating before smaller, unmyelinated (C) nociceptive neurons. This temporal difference was established from early birthdating studies based on DRG soma cell size. However, distinctions in birthdates between molecular subtypes of sensory neurons, particularly nociceptors, is unknown. Here, we assess the birthdate of lumbar DRG neurons in mice using a thymidine analog, EdU, to label developing neurons exiting mitosis combined with co-labeling of known sensory neuron markers. We find that different nociceptor subtypes are born on similar timescales, with continuous births between E9.5 to E13.5, and peak births from E10.5 to E11.5. Notably, we find that thinly myelinated Aδ-fiber nociceptors and peptidergic C-fibers are born more broadly between E10.5 and E11.5 than previously thought and that non-peptidergic C-fibers and C-LTMRs are born with a peak birth date of E11.5. Moreover, we find that the percentages of nociceptor subtypes born at a particular timepoint are the same for any given nociceptor cell type marker, indicating that intrinsic or extrinsic influences on cell type diversity are occurring similarly across developmental time. Overall, the patterns of birth still fit within the classical “two wave” description, as touch and proprioceptive fibers are born primarily at E10.5, but suggest that nociceptors have a slightly broader wave of birthdates with different nociceptor subtypes continually differentiating throughout sensory neurogenesis irrespective of myelination.

## Introduction

Sensory neurons of the dorsal root ganglion (DRG) detect somatosensory information from the skin and muscle as well as signals from visceral tissues and contain many cell types (Abraira and Ginty, 2013; Basbaum et al., 2009; Faure et al., 2020; Julius and Basbaum, 2001; Sharma et al., 2020; Usoskin et al., 2014). These sensory neurons are classically categorized based on conduction velocity which correlates with soma cell size. The largest sized soma are those of the proprioceptors, which detect muscle stretch and tension through muscle spindles and Golgi tendon organs and are the most highly myelinated. Low-threshold mechanoreceptors (LTMRs) mediating touch information, called Aβ-fibers, are the next largest myelinated fibers. Thinly myelinated Aδ-fibers, which can be LTMRs or nociceptors are the smallest myelinated fibers. Lastly, the unmyelinated C-fibers are mostly nociceptors, with one C-LTMR class of neurons.

Sensory neurogenesis begins with the delamination of neural crest cells (NCCs) from the dorsal neural tube, which starts around embryonic day 9 (E9) and continues to E13.5 in the mouse (Lallemend and Ernfors, 2012; Marmigère and Ernfors, 2007; Serbedzija et al., 1990). A variety of birthdating studies have provided evidence of two distinct waves of neurogenesis giving rise first to primarily large, myelinated neurons (proprioceptors and Aβ-LTMRs) from E9.5-E11.5, and secondly to small, unmyelinated nociceptor neurons around E10.5-E13.5 (Lawson and Biscoe, 1979; Ma et al., 1999). The birthdating pattern observed in mouse, rat, and chick suggests some overlap in the timing of the two waves, with the first preceding the second by about a day (Carr and Simpson, 1978; Kitao et al., 1996; Lawson and Biscoe, 1979; Lawson et al., 1974). Moreover, while the initial wave primarily produces large soma neurons, it is thought to include some lightly myelinated Aδ nociceptors as well. The second wave, in contrast, is thought mostly to give rise to small unmyelinated C-fiber neurons (Kitao et al., 2002; Lallemend and Ernfors, 2012; Lawson and Biscoe, 1979).

Recently, single cell RNA-sequencing data have illuminated the diversity of molecular cell types in the mouse DRG with anywhere from 11-14 distinct subtypes (Faure et al., 2020; Sharma et al., 2020; Usoskin et al., 2014). In this study, we sought to obtain more granularity as to the birthdates of different sensory populations, specifically nociceptors, using current molecular markers. We hypothesized that different nociceptor cell types might be born at different times during sensory neurogenesis. By injecting the thymidine analog 5-ethynyl-2’-deoxyuridine (EdU) at different time points during development, we permanently labeled neurons undergoing their final mitotic divisions for later identification using immunohistochemistry and RNAscope. We found that large, myelinated proprioceptors and Aβ-fibers were born at E9.5-E10.5 during the early wave consistent with previous reports. In contrast, we found that nociceptor neurogenesis occurs along a continuum from E9.5 to E13.5 with peak birthdates around E10.5-E11.5. Interestingly, we find that birth patterns of TRKA^+^ C- and Aδ-fibers are quite similar, with both being maximally born over E10.5-E11.5, while non-peptidergic C-fibers and C-LTMRs are born mostly at E11.5. Furthermore, we find that any given nociceptor subtype is born in the same percentages over this developmental window indicating that intrinsic and extrinsic factors influencing cell type occur continuously throughout this period.

## Materials and Methods

### Tissue Processing

Male and female wildtype CD1 (ICR) mice (Charles River) were crossed to each other and females were checked daily for vaginal plugs to determine date of conception. Pregnant dams were given one intraperitoneal (IP) injection of 0.5mg/ml EdU prepared in sterile 1x PBS at a dose of 10µg EdU/g animal on one day between E9.5 and E13.5. At P28, mice were anesthetized with Avertin (2,2,2-Tribromoethanol) (0.025-0.030 mL of 0.04 M Avertin in 2-methyl-2-butanol and distilled water/g mouse) and transcardially perfused, first with 0.012% w/v Heparin/PBS and then 4% paraformaldehyde (PFA) in PBS. A dorsal or ventral laminectomy exposed the spinal cord and DRGs to the fixative. The DRGs were fixed for 2 hrs at 4°C. Tissue was washed in PBS for at least one day and cryoprotected in 30% sucrose dissolved in deionized water. The right and left L2-L5 DRGs were then embedded in OCT (Tissue-Tek Optimal Cutting Temperature compound). Tissue was sectioned using a Leica CM1950 Cryostat at 30μm thickness.

### Immunohistochemistry, EdU labeling, RNAscope, and confocal imaging

Cryosections (30μm) were blocked with 1% normal goat or donkey serum/0.3% Triton X-100 (Sigma) for up to 1 hour at room temperature (RT) and incubated overnight with primary antibodies at 4°C. After washing 3 times with PBS for 10 minutes each, the appropriate secondary antibodies (Alexa 488 and 647, Invitrogen) were incubated for an hour at RT. Sections were rinsed 3 times, 10 minutes each in PBS and were permeabilized with 0.5% TritonX-100 (Sigma) for 30 minutes at RT. After washing 3 times, 5 minutes each with PBS, EdU detection solution (100µM Tris pH7.5, 4mM CuSO4, 5µM sulfo-Cy5 azide, 100mM sodium ascorbate) was applied and incubated for 30 minutes at RT protected from light. Sections were rinsed 2 times, 5 minutes each in PBS and incubated with DAPI for 1 minute. After washing 3 times in PBS sections were mounted with Aquapolymount (Polysciences Inc.), and coverslipped (Fisher). The following primary antibodies and dilutions were used: 1:500 IB4 (Invitrogen), 1:1000 rabbit anti-CGRP (Immunostar), 1:500 goat anti-TRKA (R&D Systems), 1:500 goat anti-TRKC (R&D Systems), 1:500 rabbit anti-NF200 (Sigma), 1:500 rabbit anti-TRPV1 (Alomone), 1:1000 rabbit anti-TH (Pel-Freez).

For RNAscope, tissue was collected as above, except cryosectioning was performed with 20µm sections, then stained with the RNAscope Fluorescent Multiplex Assay (Advanced Cell Diagnostics Inc., Hayward, CA). Slides were first incubated at 50°C for 1 hour. All further incubation steps were performed in a HybEZ™ II oven set to 40°C. The slides were then washed three times in 1x PBS, and incubated with Protease III (diluted 1:5) for 2.5 minutes. Slides were then washed twice with 1x PBS and hybridized with the probe for 2 hours. The slides were washed two times thoroughly using 1X wash buffer for 2 min, then incubated with Amp 1-Fl for 30 minutes. The same process (washing then treatment) was repeated for Amp 2-Fl, Amp 3-Fl and Amp 4-Fl for 15, 30 and 15 minutes, respectively. Slides were washed twice in 1x PBS, then stained for EdU, as described above starting with the 0.5% TritonX-100 step. The following probes were used with 1:50 dilution in probe diluent: *Ntrk2* (Catalog No. 423611-C2), *Mrgprd* (Catalog No. 417921-C2), and *Sst* (Catalog No. 404631-C2).

Fluorescent images were taken on a LSM880 confocal microscope with an appropriate optical slice (3 μm) with 20x objective. Images were pseudocolored using a magenta/yellow/cyan color scheme using Adobe Photoshop (Adobe) and Fiji (Schindelin et al., 2009).

### Cell counting and statistical analysis

Confocal images were counted manually in Fiji. For each experiment, three images were analyzed from each mouse, and EdU labeling was measured as a percentage of total neurons counted across all images from that mouse, or as a percentage of all neurons expressing the marker assessed in that experiment. These values were averaged for each mouse at a given timepoint, with a minimum of three mixed-gender mice per timepoint (see Table S6 for exact n), and the average ± SEM is reported in the figures. Statistical analysis for each experiment was performed by one-way ANOVA with a post-hoc Tukey test. F statistics and *p* values for the ANOVA are reported in the text, and p values for pairwise comparisons of each time point are reported in the supplemental tables.

## Results

### Larger myelinated DRG neurons are born before smaller unmyelinated neurons

To investigate the birthdates of different sensory populations, we injected EdU into pregnant dams at a range of time points from E9.5 to E13.5. EdU is incorporated into nuclear DNA during S-phase, and thereby permanently labels all cells undergoing their final mitotic divisions on the day of injection. We then harvested DRGs from P28 mice, when the sensory neurons have matured (Lallemend and Ernfors, 2012). We first quantitated the number of EdU^+^ neurons as a percentage of total DRG neurons to measure changes in birth rate over time. We found that neuronal births in the DRG occur continuously from E9.5 to E13.5, with significantly greater number of neurons born on E10.5 and E11.5, as determined by one-way ANOVA (*F*(4, 15) = 35.12, *p* < 0.0001) with post-hoc Tukey tests (*p* = 0.0003 or smaller comparing these time points to the later time points, Fig. 1A and Table S1). Curiously, while the overall neuronal birth range is consistent with prior studies, we see the greatest number of neurons born one day earlier than previously reported for mouse lumbar DRGs (Lawson and Biscoe, 1979).

**Figure 1.**
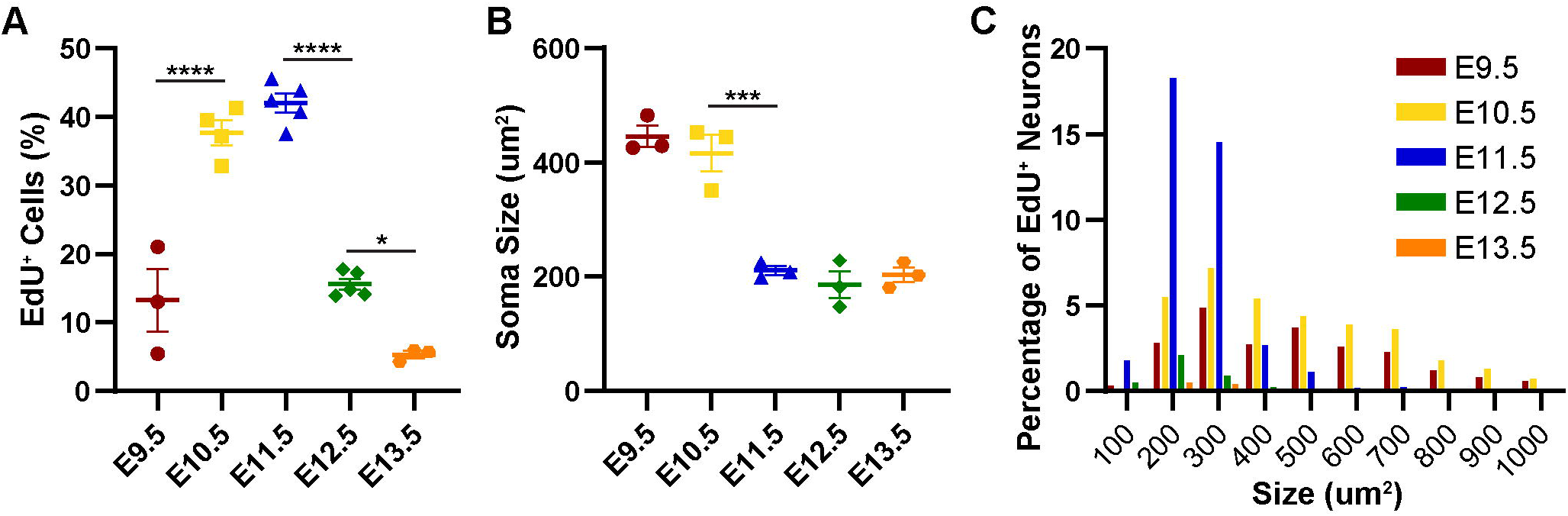
Sensory neurons born early in DRG development are larger than those born later. (A) Most sensory neurons undergo their final mitotic divisions and are born between E9.5 – E13.5, with a significant increase in birth rate at E10.5 – E11.5, as determined by one-way ANOVA (*F*(4, 15) = 35.12, *p* < 0.0001). (B) Measured at P28, the soma of sensory neurons born at E9.5 – E10.5 are significantly larger than those born later, as determined by one-way ANOVA (*F*(4, 10) = 37.62, *p* < 0.0001). (C) Histogram showing numbers of sensory neurons born each day as a percentage of total EdU^+^ neurons, binned by size. Neurons born at E9.5 and E10.5 span the entire range from 100 – 1000 µm^2^, while those born at E11.5 and beyond are mostly in the 100 – 500 µm^2^ range. Asterisks indicate significant changes identified with a post-hoc Tukey’s test; see Supplemental Table 1 for multiple all statistics.

We next assessed the sizes of EdU^+^ neurons labeled at different time points by measuring the cross-sectional area of cell somas. We found that neurons born earlier (E9.5 and E10.5) were larger, averaging slightly over 400µm^2^, while those born from E11.5—E13.5 were significantly smaller, at around 200µm^2^ (Fig. 1B). Moreover, we found that while smaller neurons are born continuously from E9.5—E13.5, with most being born at E11.5, the largest neurons (greater than 500µm^2^) are born almost exclusively on E9.5 and E10.5 (Fig. 1C). These findings are consistent with previous reports of cell size by birthdate (Kitao et al., 2002; Lawson and Biscoe, 1979).

### TRKA^+^ neurons, both myelinated and unmyelinated, are born at similar times

We next wanted to examine specific sensory populations to determine whether there are differences in their birth dates. We were most interested in the wide array of nociceptive neurons, and whether there exists any temporal difference in birth day for different subtypes. We started by assessing TRKA^+^ (tropomyosin receptor kinase A) nociceptors, which we classified as Aδ-fibers if they co-expressed the myelination marker NF200 (neurofilament 200), or C-fibers if they did not (Fig. 2A) (Bachy et al., 2011; Sharma et al., 2020; Usoskin et al., 2014). As a whole, TRKA^+^ neurons are born on a continuum from E9.5 to E13.5, with a significantly greater number of births as a percentage of the whole DRG at E10.5 and E11.5 (Fig. 2B, *F*(4, 15) = 23.83, *p* < 0.0001; see Table S2 for all statistics). The majority of TRKA^+^ neurons are born during this 2-day window, with 40% born on E10.5, and another 37% on E11.5 (Fig. 2C, *F*(4, 15) = 11.33, *p* = 0.0002).

**Figure 2.**
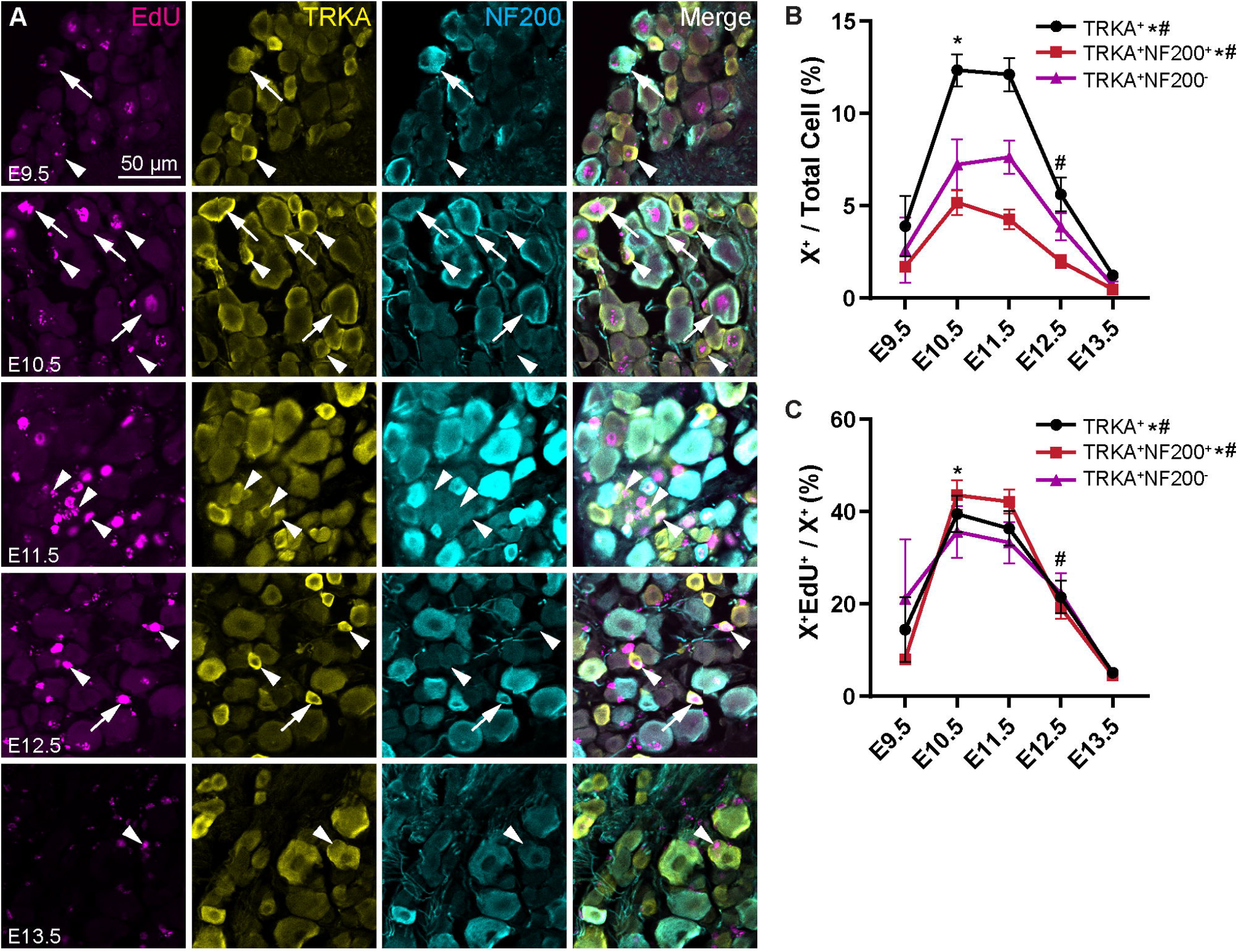
TRKA^+^ Aδ- and C-fibers are born in similar numbers over developmental time. (A) Representative images of immunohistochemistry showing EdU, TRKA, and NF200 at P28 following injection of EdU at the indicated timepoints from E9.5 – E13.5. Arrows show triple-positive EdU^+^ TRKA^+^ NF200^+^ cells (EdU^+^ Aδ-fibers); arrowheads show EdU^+^ TRKA^+^ NF200^-^ cells (EdU^+^ C-fibers). (B, C) Quantitation of total TRKA^+^ Aδ-fibers, and TRKA^+^ C-fibers labeled by EdU at each time point as a percentage of all DRG neurons (B) or that specific neuronal type (C). Overall, births of the TRKA^+^ fibers are maximal from E10.5—E11.5; this trend holds for both the Aδ- and TRKA^+^ C-fibers. * indicates a significant increase in births from E9.5 to E10.5, # indicates a significant decrease in births from E10.5 to E11.5; see Supplemental Table 2 for all statistics.

Given that myelinated fibers as a whole are known to develop earlier during neurogenesis, we hypothesized that TRKA^+^ neurons born earlier were likely to be Aδ-fibers, while those born later were likely to be C-fibers. Surprisingly, we find that there was not a significant difference in births for either of the TRKA^+^ Aδ- or C-fiber populations from E10.5 to E11.5 (Fig. 2B). During this window, about 85% of TRKA^+^ Aδs were born, and 69% of TRKA^+^ C-fibers, with each split about evenly across the two days (Fig. 2C). Thus, it appears that TRKA^+^ nociceptors are born primarily at E10.5—E11.5, regardless of whether or not they are myelinated.

### Births of nociceptors of any given subtype are similar across developmental time

We next assessed the birth rates of other subtypes of nociceptors, starting with non-peptidergic, IB4^+^ C-fibers (Fig. 4A). Given that these are another population of small, unmyelinated neurons, we expected them to be born relatively late in development. We found that IB4^+^ fibers showed a significant peak birth date at E11.5 (Fig. 3D, *F*(4, 15) = 50.31, *p* < 0.0001) with 50% of the population born that day (Fig. 3E *F*(4, 15) = 31.94, *p* < 0.0001; See Table S3 for all statistics). C-low threshold mechanoreceptors (C-LTMRs), marked by tyrosine hydroxylase (TH, Fig. 3B), similarly show a peak birth date at E11.5 (Fig. 3D, *F*(3, 8) = 9.559, *p* = 0.0051), again with 50% of TH^+^ neurons born that day (Fig. 3E, *F*(3, 8) = 13.31, *p* = 0.0018, see table S3).

**Figure 3.**
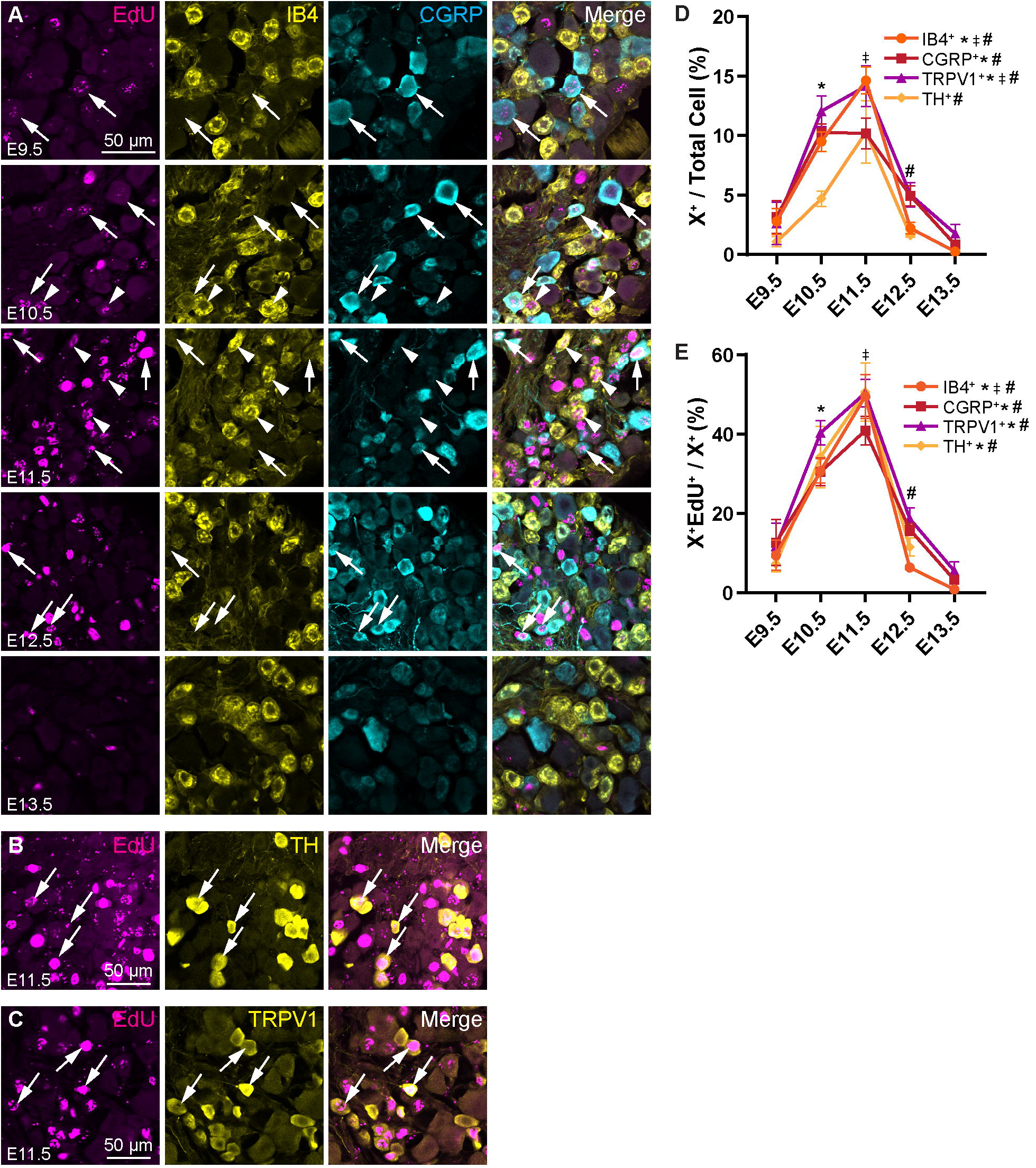
Non-peptidergic C-fibers and C-LTMRs are born mostly at E11.5, but significant portions of the populations are also born at E10.5. (A) Representative images showing EdU, IB4, and CGRP immunohistochemistry at P28 following injection of EdU at the indicated timepoints from E9.5 – E13.5. Arrows show EdU^+^ CGRP^+^ peptidergic nociceptors, arrowheads show EdU^+^ IB4^+^ non-peptidergic nociceptors. (B, C) Representative images of EdU^+^ TH^+^ (B) and EdU^+^ TRPV1^+^ (C) cells labeled at E11.5. (D, E) Quantitation of nociceptor populations labeled by EdU at each time point as a percentage of all DRG neurons (D) or that specific nociceptor subtype (E). Peptidergic nociceptors appear to be born continuously across developmental time with a peak spanning E10.5—E11.5, while non-peptidergic C-fibers and C-LTMRs have peak birth rates more specifically at 11.5. * indicates a significant increase in births from E9.5 to E10.5, ∼ indicates a significant increase in births from E10.5 to E11.5, # indicates a significant decrease in births from E11.5 to E12.5; see Supplemental Table 3 for all statistics.

**Figure 4.**
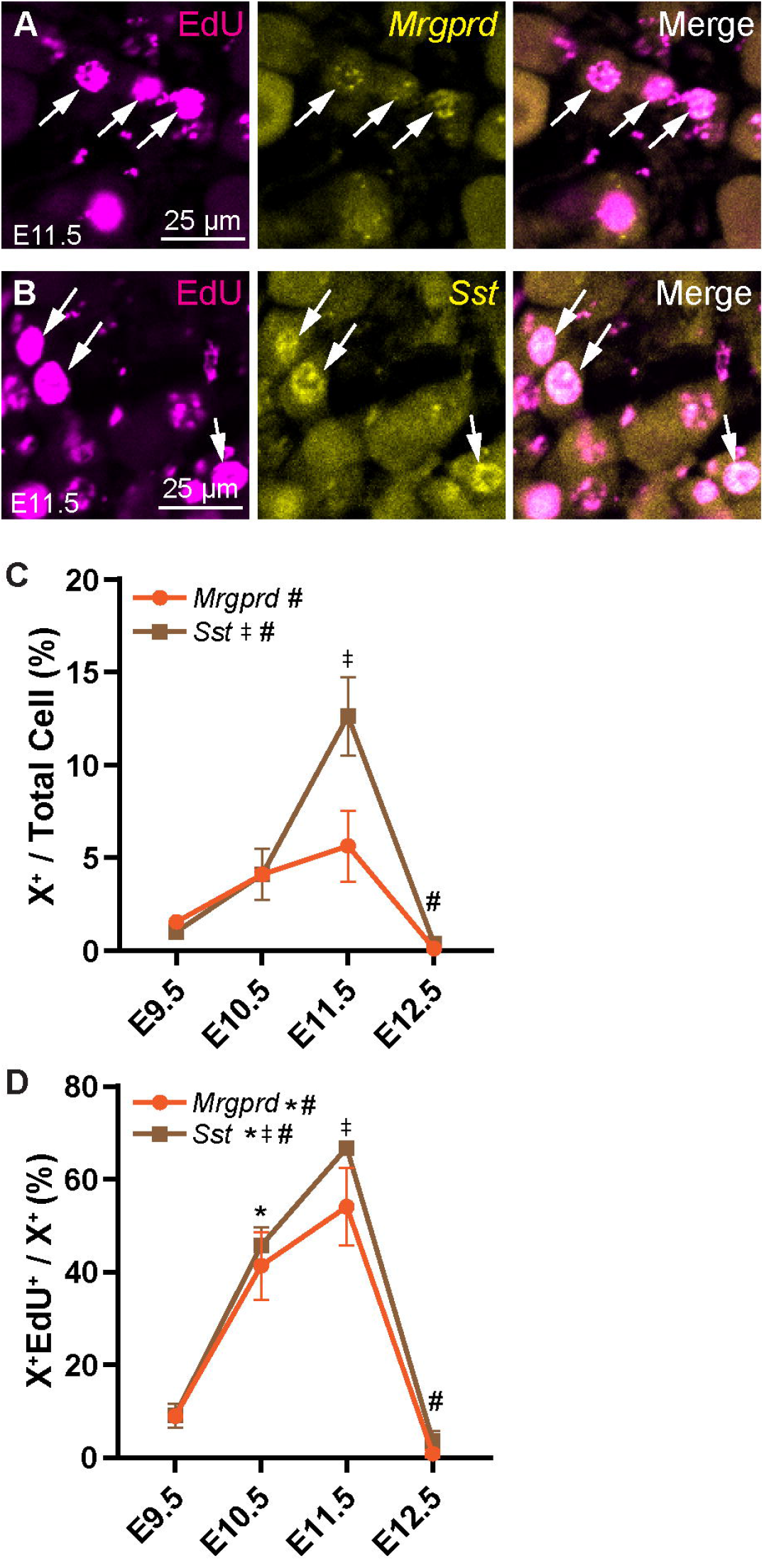
Birth rates in nociceptor populations identified by RNAscope are similar to those identified with antibody staining. (A, B) Representative images of EdU labeling following RNAscope for *Sst* (A) and *Mrgprd* (B) after EdU injection at E11.5. (C, D) Quantitation of *Sst+* and *Mrgprd*^+^ labeled by EdU at each time point as a percentage of all DRG neurons (C) or those expressing that marker (D). ∼ indicates a significant increase in births from E9.5 to E10.5, * indicates a significant increase in births form E10.5 to E11.5, # indicates a significant decrease in births from E11.5 to E12.5; See Supplemental Table 4 for all statistics.

Next, we examined the birth date of peptidergic nociceptors expressing calcitonin gene-related peptide (CGRP, Fig. 3A) and nociceptors expressing the capsaicin receptor, transient receptor potential cation channel subfamily V member 1 (TRPV1, Fig. 3C). Like TRKA, CGRP is expected to label both peptidergic C- and Aδ-fibers (Sharma et al., 2020; Usoskin et al., 2014). Therefore, similar to the TRKA^+^ nociceptors, CGRP^+^ neurons have a broader peak distribution than the non-peptidergic IB4^+^ nociceptors and C-LTMRs with a significant increase in birth at E10.5 that remains elevated through E11.5 (Fig. 3D, *F*(4, 15) = 15.51, *p* < 0.0001). About 30% of CGRP^+^ neurons are born on E10.5, with another 40% the following day (Fig. 3E, *F*(4, 15) = 20.38, *p* < 0.0001). Expression of TRPV1 is also expected to broadly overlap with CGRP (Cavanaugh et al., 2011; Lallemend and Ernfors, 2012; Sharma et al., 2020; Usoskin et al., 2014). Indeed, we find birth dates of TRPV1^+^ neurons to be similar to those of the CGRP^+^ and TRKA^+^ population, with a wider peak of cell birth spanning E10.5—E11.5 (Fig. 3D, *F*(4, 14) = 15.16, *p* < 0.0001). The majority of the TRPV1^+^ population is born during this timeframe, with 40% born on E10.5 and 50% on E11.5 (Fig. 3E, *F*(4, 14) = 29.64, *p* < 0.0001).

We also used RNAscope to investigate birth dates of other neuronal populations. *Mrgprd* (MAS related GPR family member D, Fig. 4A) is expressed in a subpopulation of IB4^+^ polymodal nociceptors (Lallemend and Ernfors, 2012; Usoskin et al., 2014). While births of *Mrgprd*^*+*^ neurons appear to increase until E11.5 before steeply dropping, only the difference between E11.5 and E12.5 was found to be significant, suggesting that the overall birth rate of this population is relatively constant across developmental time (Fig. 4C, *F*(3, 8) = 6.454, *p* = 0.0157; see Table S4 for all statistics). Still, most *Mrgprd*^*+*^ neurons are born from E10.5—E11.5 (Fig. 4D, *F*(3, 8) = 20.63, *p* = 0.0004). Somatostatin (*Sst*) marks a major population of non-peptidergic itch-sensitive neurons, or pruriceptors (Fig. 4B). Like the IB4^+^ and *Mrgprd*^*+*^ non-peptidergic fibers, most of the *Sst*^+^ population is born at E11.5 (Fig. 4C, *F*(3, 8) = 19.52, *p* = 0.0005), with 67% of *Sst*^+^ neurons born that day (Fig. 4D, *F*(3,8) = 124.8, *p* < 0.0001).

In summary, we report that nociceptors, both myelinated and unmyelinated, are primarily born from E10.5—E11.5, with a slight preference for non-peptidergic C-fibers to be born in the latter half of this window. Notably, we find that all nociceptors are born in similar percentages across developmental time (Fig. 3E and Fig. 4D). For example, approximately 31% of the IB4^+^, 34% of the TH^+^, and 40% of the TRPV1^+^ population are born at E10.5 (Fig. 3E). This suggests that intrinsic or extrinsic factors are equivalently affecting diverse nociceptor populations during neurogenesis.

### Large touch and proprioceptive fibers are born predominantly at E10.5

Finally, we looked at the larger myelinated fibers, including LTMR TRKB^+^ Aβ-fibers (Fig. 5A with RNAscope for *Ntrk2*), and touch and proprioceptive TRKC^+^ fibers (Fig. 5B). We found that the myelinated fibers as a whole (NF200^+^), and *Ntrk2*^+^ and TRKC^+^ populations specifically, had a significant peak in neuronal birth at E10.5, with a sharp decline one day later (Fig. 5C; NF200 *F*(4, 15) = 16.00, *p* < 0.0001; *Ntrk2 F*(3, 8) = 10.16, *p* = 0.0042; TRKC *F*(3, 8) = 6.653, *p* = 0.0145; see Table S5 for all statistics). Also, the greatest percentage of each of these population was born on E10.5: 41% of NF200^+^ neurons, 54% of *Ntrk2*^*+*^ neurons, and 32% of TRKC^+^ neurons (Fig. 2D; NF200 *F*(4, 15) = 21.58, *p* < 0.0001; *Ntrk2 F*(3, 8) = 40.16, *p* < 0.0001; TRKC *F*(3, 8) = 10.69, *p* = 0.0036). Overall, we find that births of large, myelinated fibers appear to be restricted to early neurogenesis, from E9.5—E10.5, compared to the nociceptors which are born more continuously throughout development.

**Figure 5.**
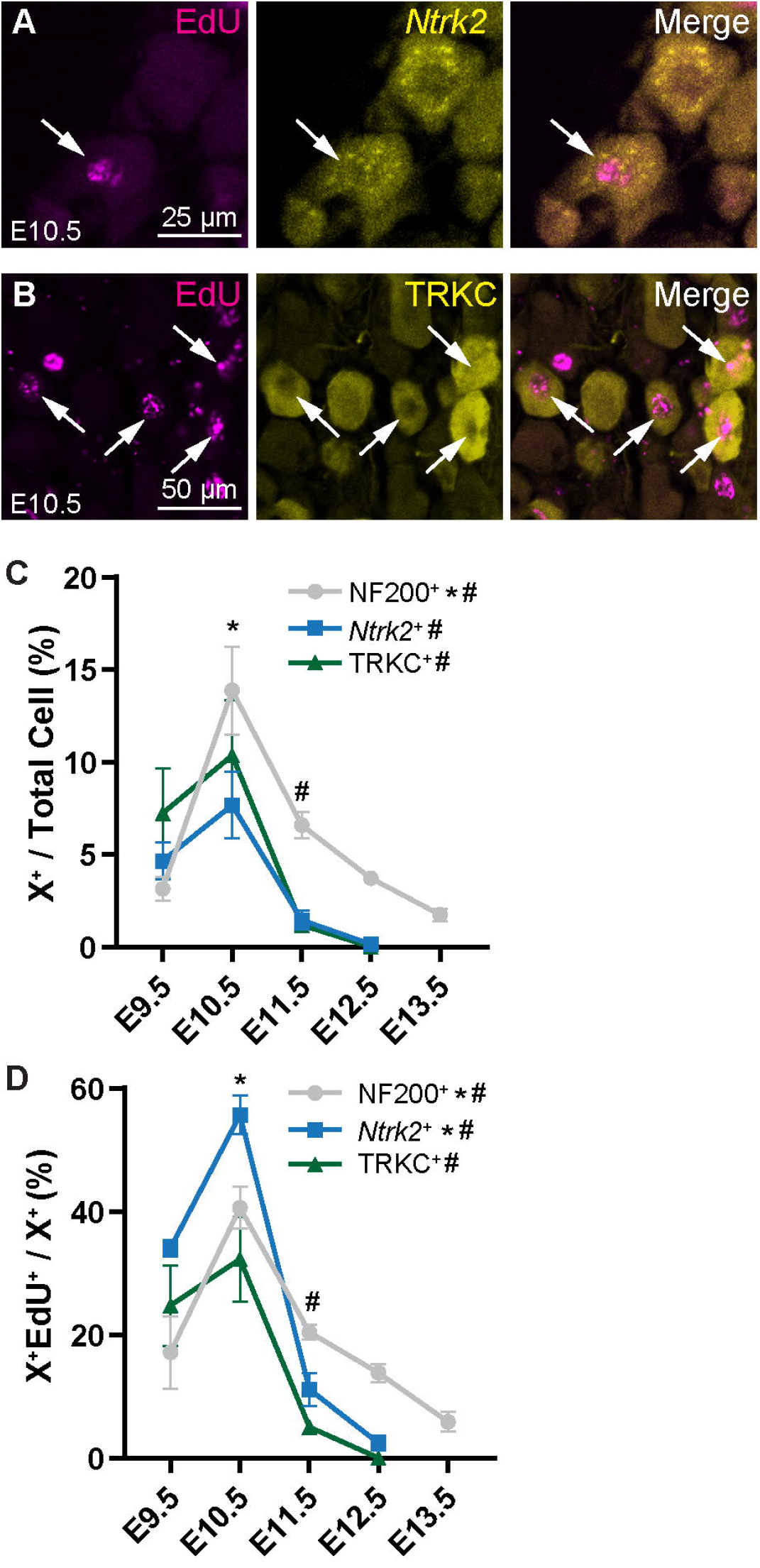
LTMRs and proprioceptive neurons are born primarily at E10.5. (A) Representative images of EdU^+^ *Ntrk2*^*+*^ (detected by RNAscope) cells labeled at E10.5. (B) Representative images of EdU^+^ TRKC^+^ cells labeled at E10.5. (C, D) Quantitation of NF200^+^ (see Fig. 2), *Ntrk2*^*+*^, and TRKC^+^ neurons born at each time point as a percentage of all DRGs neurons (C) or all neurons of that subtype (D). Compared to nociceptors, these populations show a peak birth at E10.5. * indicates a significant increase in births from E9.5 to E10.5, # indicates a significant decrease in births from E10.5 to E11.5; see Supplemental Table 5 for all statistics.

## Discussion

In this study, we explore the birth patterns of a variety of sensory neuron types using the thymidine analog EdU to permanently label them on the day they become post-mitotic, their birth day. We predicted that we would see the previously reported two waves of neurogenesis, differentiated by cell size, but that we might also find differences in the timing of birth of various nociceptor populations. Surprisingly, we found that while the largest neurons, *Ntrk2*^*+*^ and TRKC^+^ myelinated touch and proprioceptive neurons, are indeed born earliest, the smaller Aδ population was born on a similar timescale as peptidergic C-fibers, with nonpeptidergic C-fibers and C-LTMRs born last (summarized in Fig. 6B). We make these conclusions primarily based on the peak birthdate, assessed as the percentage of newly-born neurons of a given subtype over total DRG neurons (Figures 2B, 3D, 4C, 5C).

**Figure 6.**
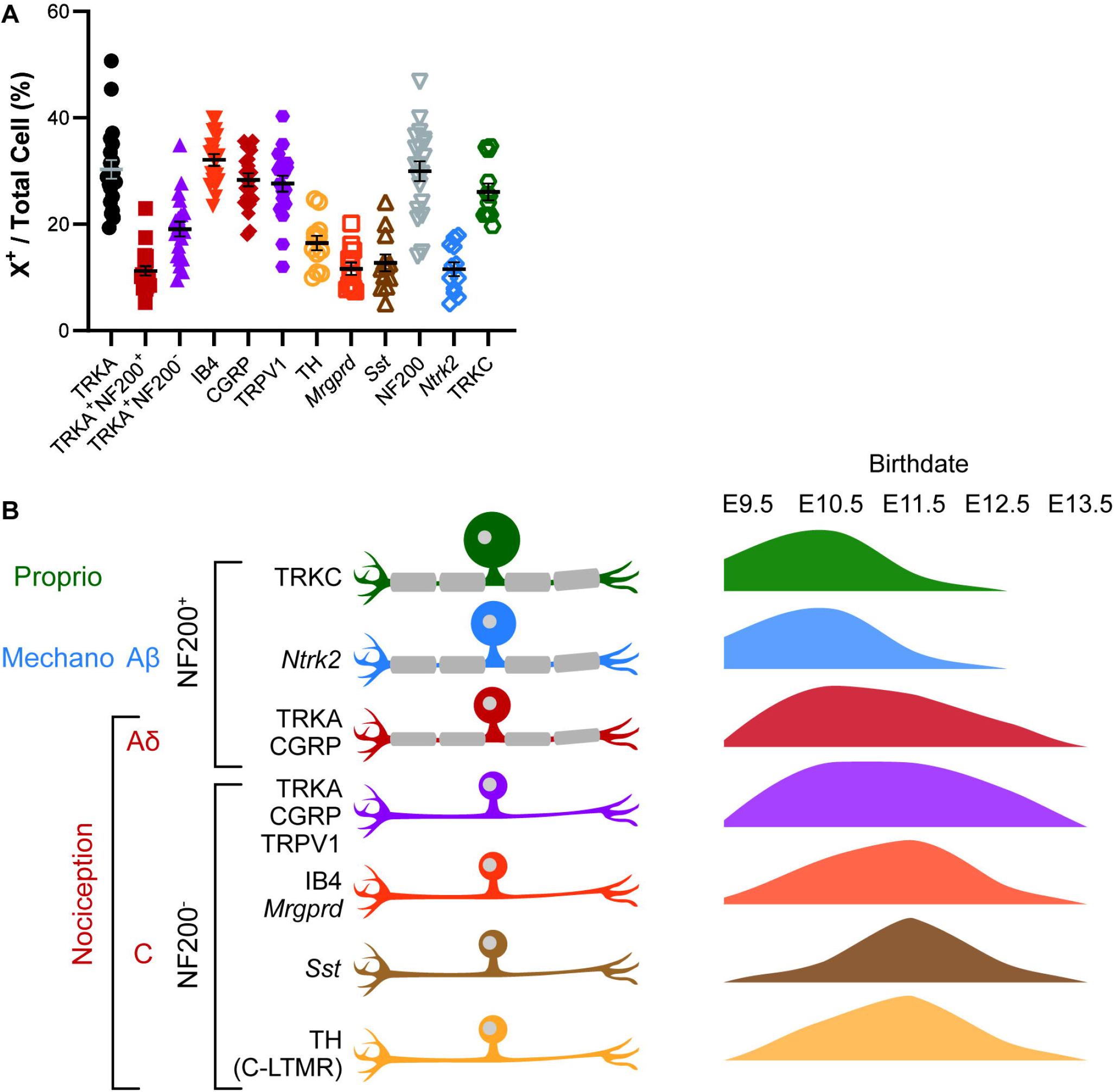
Average percentages of each cell type and summary of neurogenesis by cell type. (A) Quantitation of average percentages for each cell type. Each datapoint is the average percentages from 3 lumbar DRG sections of a single mouse. See Table S6 for n values. (B) Summary of neurogenesis with cell populations and relevant markers investigated in this study shown on the left, and graphical representations of their birthdates on the right. Proprioceptive TRKC^+^ and mechanosensitive *Ntrk2*^*+*^ Aβ-fibers are born early, peaking at E10.5, while nociceptive neurons, including Aδ- and C-fibers are born more continuously with peaks around E10.5—E11.5.

As a whole, smaller diameter nociceptive neurons are born more continuously across developmental time than the larger diameter neurons (Fig. 1C). Notably, we find that the birth rate of different nociceptive populations is similar, with substantial portions of each subtype born at both E10.5 and E11.5 (Figures 2C, 3E, 4D). This suggests that intrinsic and extrinsic factors leading to the diversification of nociceptors in the mature DRG occurs similarly over developmental time. Exactly how factors influencing cell type exert their effects at the progenitor stage, post-mitotically, or both is an open question (Faure et al., 2020; Sharma et al., 2020).

One possible confounding factor in our study is the exact differences in timing of cell cycle exit that can be resolved with EdU labeling. This is dependent on both the pharmacokinetics of EdU clearance from the animal, and the rate of proliferation of neuronal precursors. To the best of our knowledge, timing of EdU clearance has not been well investigated, and reports of doubling times of chick or sheep NCCs range from 5—26 hours (Maxwell, 1976; Ridenour et al., 2014; Zeuner et al., 2018). If the longer end of the range is more accurate, then it is conceivable that a cell labeled by EdU on one day may undergo one final round of proliferation and exit the cell cycle the following day. This could result in “spillover” where the detected birth day of a given neuron is off by up to one day. However, the sharp increases and decreases we see in numbers of EdU-labeled cells between days suggest that EdU does indeed mark terminal differentiation within a 24-hour period. Furthermore, the percentage born at a given time point for a given marker should add up to 100% for all time points if EdU was accurately marking neurons born just on that day. We found that for most of the markers used in our study, the sum was within ± 5% of 100%, with the exception of TRKC (62%), TRKA (117%), *Sst* (125%), and TRPV1 (127%). Part of the deviation from summing to 100% can be attributed to biological noise and inherent error in counting a limited number of sections/DRG. As can be seen from summary data of the average percentages for any given cell type marker in the lumbar DRG, the percentage for any given marker counted from 3 DRG sections can vary widely (Fig. 6A). Even with these possible errors, the fact that counts from multiple animals over multiple birth dates adds close to 100% indicates that there is not much spillover of the EdU marking neurons a day later.

It also bears noting that approximately 5% of DRG sensory neurons originate from neural crest boundary cap cells as part of a “third wave” of neurogenesis beginning by E11.5 (Marmigère and Ernfors, 2007; Maro et al., 2004). At least 87% of these cells become nociceptors, split between IB4^+^ and CGRP^+^ neurons, with almost none producing larger diameter cells (Maro et al., 2004). These contributions to the overall pool of DRG neurons, while small, likely contribute to the maximal birth rates of nociceptive neurons seen at E11.5.

Overall, the data we present here help to elucidate the temporal differences in birth rates of various sensory neuronal populations. Previous reports delineated patterns of birth primarily on the basis of cell size. We expand on this and show that nociceptor subtypes are born at similar rates, regardless of whether they are Aδ- or C-fiber nociceptors. These findings will be helpful in further studies of the earliest developmental processes in nociceptor neurons.

## Figure Legends

**Supplemental Table 1**. Statistical data pertaining to Figure 1.

**Supplemental Table 2**. Statistical data pertaining to Figure 2.

**Supplemental Table 3**. Statistical data pertaining to Figure 3.

**Supplemental Table 4**. Statistical data pertaining to Figure 4.

**Supplemental Table 5**. Statistical data pertaining to Figure 5.

**Supplemental Table 6**. Number of biological replicates for each experiment.

## Funding

This work was supported by F31 NS111796 and the William F. and Grace H. Kirkpatrick Award to M.A.L. and the Rita Allen Foundation Award in Pain, Welch Foundation I-1999-20190330, President’s Research Council Award, the Kent Waldrep Foundation, and NIH R01 NS100741 to H.C.L.

## Acknowledgements

We thank Tou Yia Vue for help with EdU staining, and Marghi Jani for help with tissue collection. We thank Jane Johnson for critical reading of the manuscript.

## Declaration of Interests

The authors declare no competing interests.

## Notes

### Competing Interest Statement

The authors have declared no competing interest.

